# Structural models of full-length JAK2 kinase

**DOI:** 10.1101/727727

**Authors:** Pelin Ayaz, Henrik M. Hammarén, Juuli Raivola, Dina Sharon, Stevan R. Hubbard, Olli Silvennoinen, Yibing Shan, David E. Shaw

**Affiliations:** D. E. Shaw Research, New York, NY 10036, USA; Faculty of Medicine and Life Sciences, University of Tampere, Tampere, Finland; Kimmel Center for Biology and Medicine, Skirball Institute and Department of Biochemistry and Molecular Pharmacology, New York University School of Medicine, New York, NY 10016, USA; Fimlab Laboratories, Pirkanmaa Hospital District, Tampere, Finland; Institute of Biotechnology, University of Helsinki, Helsinki, Finland; Department of Biochemistry and Molecular Biophysics, Columbia University, New York, NY 10032, USA

## Abstract

The protein JAK2 is a prototypical member of the Janus kinase family, and mediates signals from numerous cytokine receptors. The constitutively active V617F mutant of JAK2 is prevalent in many bone marrow disorders, blood cancers, and autoimmune diseases, and is an important drug target. Structures have been determined for each of the four individual domains making up JAK2, and for certain pairs of these domains, but no structure of full-length JAK2 is available, and thus the mechanisms underlying JAK2 regulation and the aberrant activity of the V617F mutant have been incompletely understood. Here we propose structural models of full-length JAK2 in both its active and inactive forms. Construction of these models was informed by long-timescale molecular dynamics simulations. Subsequent mutagenesis experiments showed that mutations at the putative interdomain interfaces modulated JAK2 activity. The models provide a structural basis for understanding JAK2 autoinhibition and activation, and suggest that the constitutive activity of the V617F mutant may arise from a dual effect of destabilizing the inactive conformation and stabilizing the active conformation.

## Introduction

Members of the Janus kinase (JAK) family of non-receptor tyrosine kinases act immediately downstream of various cytokine receptors that mediate the signaling of interleukins, interferons, and hormones (such as growth hormone, erythropoietin, and leptin).^1^ The four mammalian JAKs (JAK1, JAK2, JAK3, and TYK2) are constitutively bound to the intracellular segments of cytokine receptors^2^ and phosphorylate signal transducers and activators of transcription (STATs),^3^ thereby regulating the immune system, hematopoiesis, cellular metabolism, and growth.^4^ Many mutations in JAK proteins lead to cytokine-independent signaling, and underlie a number of severe human diseases, including various immune-related conditions and certain forms of cancer.^5, 6^ The mutation V617F in JAK2 is the most prevalent, and is strongly associated with myeloproliferative neoplasms, where it is found in nearly all patients with polycythemia vera, and about half of all patients with essential thrombocythemia or idiopathic myelofibrosis.^7, 8^ As a result, the V617F mutant of JAK2 is an important drug target.^9^ The structural basis of JAK2 regulation and the aberrant activity of V617F, however, are incompletely understood.

Like other JAKs, JAK2 consists of four domains. From the N terminus to the C terminus, these are the 4.1 protein ezrin/radixin/moesin (FERM) domain, the Src homology-2 (SH2)–like domain, the Janus homology-2 (JH2) pseudokinase domain, and the Janus homology-1 (JH1) tyrosine kinase domain. The FERM and SH2 domains bind constitutively to the intracellular regions of cytokine receptors;^10^ these two domains have been structurally resolved in complex with a cytokine receptor peptide,^11, 12^ and can be considered a single structural unit of JAK2. The FERM domain is composed of subdomains F1, F2, and F3; subdomains F1 and F3 interact with the SH2 domain, and F2 is thought to interact with the cell membrane.^11^ When a JAK is in its inactive state, the JH2 and JH1 domains have been suggested to adopt a stable complex structure in which JH2 autoinhibits JH1.^13–15, 17^ This JH2-JH1 complex can thus be considered another structural unit in the inactive state.^16, 17^ A study in insect cells found that a JH2-JH1 construct from JAK2 was incompletely autoinhibited,^18^ suggesting that interactions between the JH2-JH1 unit and the FERM-SH2 unit are integral to complete autoinhibition of the full-length JAK2 protein, but no full-length structure of JAK2 is available.

Electron microscopy experiments have revealed distinct (compact and elongated) conformational states of JAK1,^19^ suggesting that monomeric JAKs exist in dynamic conformational equilibrium. The compact conformation is likely to accommodate more extensive interaction between the two structural units and thus reflect the autoinhibited state of JAK1. Cytokines induce the formation of active cytokine receptor dimers,^20^ which in turn may mediate the dimerization and activation of JAK2.^21^ It has been proposed that the dimerization releases the JH1 kinase domains from the autoinhibitory monomer structure, allowing trans-phosphorylation of the two JH1 domains at the Y1007 and Y1008 residues in the activation loop.^22^ It has also been suggested that the JH2 domain is likely to be crucial to the dimerization, and that JAK2 mutations such as F595A,^23, 24^ F739R,^25^ and others that abrogate ATP binding to JH2^26^ weaken the constitutively high kinase activity of the V617F mutant, possibly by undermining the structural integrity of the JH2 domain and thus hindering JAK2 dimerization.

Many pathogenic JAK2 mutations cluster in the SH2-JH2 linker (exon 12) and in the vicinity of the V617F residue (exon 14). For both JAK2 and TYK2, the resolved structures of the inactive JH2-JH1 units have shown that the SH2-JH2 linker is involved in the JH2-JH1 interaction; the V617F mutation site interacts directly with this linker, and it has been speculated that the mutation destabilizes the (inactive) JH2-JH1 interaction.^16^ Cell-transfection experiments, moreover, have indicated that the mutation enhances (active) JAK2 dimerization.^27^ A deeper mechanistic understanding of the constitutive activity of V617F, however, has been unattainable without structures for full-length inactive and active JAK2.

Here we propose models of full-length active and inactive JAK2. We first used long-timescale molecular dynamics (MD) simulations to inform the development of a structural model of inactive JAK2, by assembling our previous model of the JH2-JH1 unit^16^ and a resolved structure of the FERM-SH2 unit.^11^ The FERM-SH2 unit reinforces the autoinhibition of JH1 by JH2 in the resulting model. In further simulations, conformational changes in the inactive model suggested an active JAK2 monomer in which JH1 is released from direct and specific interactions with the other domains. Based on this active monomer model, we constructed a model of an active JAK2 dimer. The V617F mutation is involved in FERM-JH2 interactions in both the inactive and active models, potentially destabilizing the inactive state and stabilizing the active state. To test our models, we performed mutagenesis experiments on the V617F mutant, and found that mutations predicted to destabilize the inactive state indeed increased V617F activity, and that mutations predicted to destabilize the active state suppressed V617F activity. Together, our full-length structural models suggest architectures that may underpin JAK2 regulation and signaling, and offer a structural explanation for the constitutive activity of the V617F mutant.

## Results

### A structural model of full-length autoinhibited JAK2

We first set out to construct a structural model of full-length JAK2 based on the two resolved two-domain structures, the JH2-JH1 structure^16, 17^ and the FERM-SH2 structure.^11^ We conjectured that, while FERM is interacting primarily with cytokine receptor and cell membrane,^22^ SH2 may join JH2 in interacting with JH1 to potentiate complete autoinhibition. We simulated the association of JH1 and SH2 without any prior assumption of the binding pose, aiming to generate a structural model for SH2 interaction with JH1 in the autoinhibition. Taking the approach we used to generate the JH2-JH1 model,^16^ we selected structural models generated by the MD simulations according to analyses that took into account existing functional data on JAKs.

We initially placed the structures of SH2 and JH1 in separation from one another with an arbitrary relative orientation in an explicit solvent box (Figure 1A, state 1), and launched 20 independent MD simulations, each 3 μs in length, from this initial simulation system. These simulations, in which SH2 and JH1 developed sustained contact with one another that persisted for the duration, produced 20 preliminary structural models of an SH2-JH1 complex (Figure 1A, state 2). Based on the experimentally resolved FERM-SH2 and JH2-JH1 structures, we then extended each of these SH2-JH1 models to a preliminary full-length structure by placing FERM into its position relative to SH2 and JH2 into its position relative to JH1. From these 20 preliminary models, we first excluded the eight that contained steric clashes between FERM-SH2 and JH2 and/or between JH2-JH1 and FERM. We then excluded another eight models in which either the cytokine receptor-binding interface of the FERM-SH2 unit was occupied by the JH2-JH1 unit, or the four domains were assembled into highly elongated structures that did not appear to reinforce the autoinhibition of JH1 by JH2. Another two models were excluded due to unlikely interactions between the JH2-JH1 unit and the cytokine receptor peptide bound to the FERM-SH2 unit. From the remaining two models, we selected the one that allowed for more extensive membrane interaction and in which the V617F mutation site was involved in interdomain interactions (Figure 1A, state 3). We also considered the possibility of SH2 interacting directly with JH2 in the inactive structure of JAK2 and simulated the association of these two domains; we derived an alternative full-length model (Figure S1F) from this SH2-JH2 model, but discarded the full-length model because it did not remain stable in simulations.

**Figure 1:**
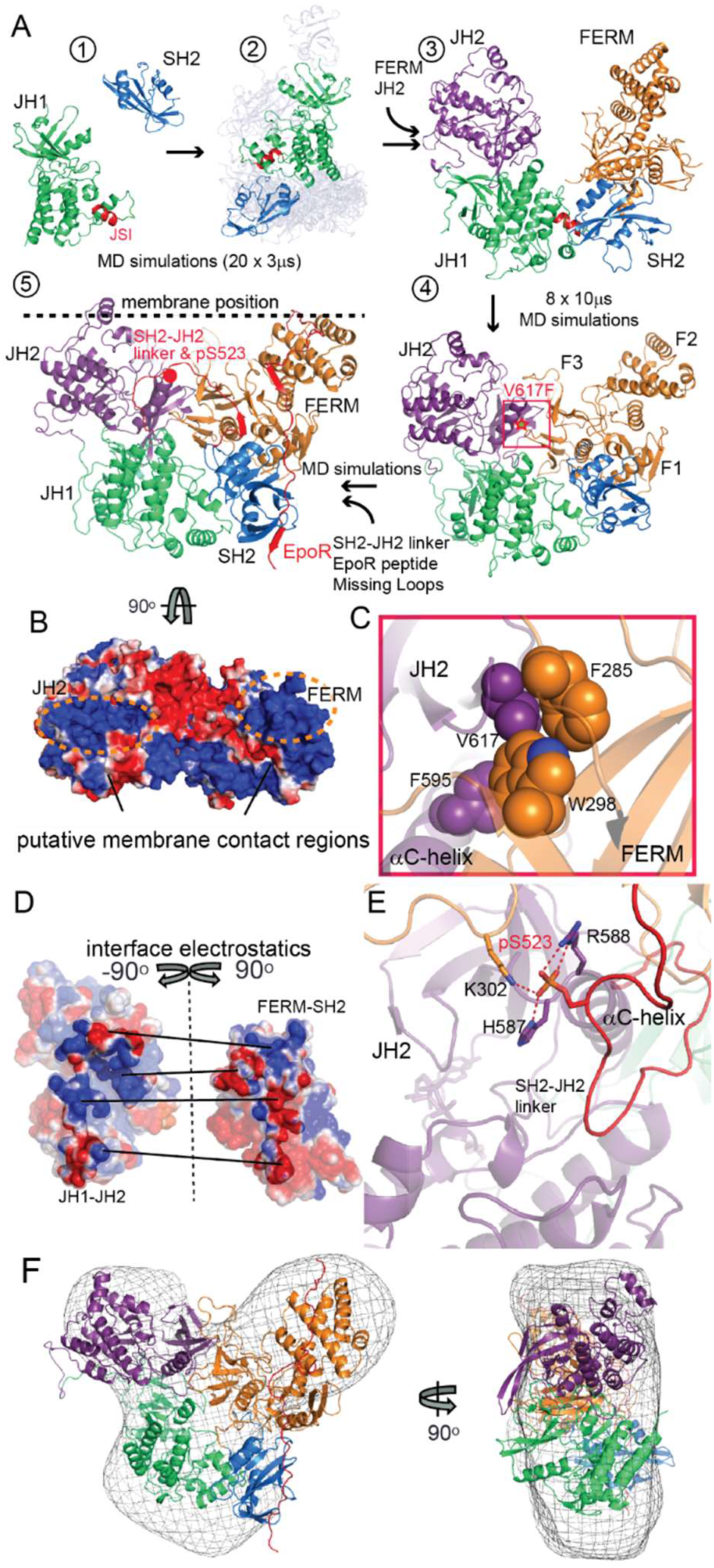
The inactive monomer model. (A) Generation of the model: (1) simulations of SH2 (blue) associating with JH1 (green); (2) the SH2-JH1 docking poses from the last frames of the 20 SH2-JH1 simulations, with the selected pose shown in solid colors; (3) the four-domain model obtained by adding FERM (orange) and JH2 (purple) to the selected SH2-JH1 pose; (4) the four-domain model obtained by simulating the model shown in (3), with the V617 region highlighted by a red square (a close-up of this region is shown in panel C); (5) the inactive monomer model of JAK2 with the SH2-JH2 linker loop and EpoR peptide added in red (see Figure S1A for the loop modeling) and a dashed line indicating the putative position of the cell membrane. (B) Surface electrostatics of the juxtamembrane region. (C) The V617 region. V617 is part of a hydrophobic cluster at the FERM-JH2 interface. (D) Electrostatic properties of the JH2-JH1 and FERM-SH2 units at their mutual interface. Straight lines connect the regions in contact at the interface. (E) Close-up of the phosphorylated S523 residue. (F) Fit of the inactive JAK2 model into a JAK1 EM envelope^19^ (front and side views).

In the selected model, SH2 interacts with the JAK-specific insertion (JSI) helix of JH1, a helix which is unique to JAK kinases,^28^ and the conserved positively charged, putative membrane-interaction patch in the FERM F2 subdomain^11^ is aligned with positively-charged patches on JH2, such that both patches can favorably interact with the membrane surface (Figure 1B and Figure S1A). We carried out eight 10-µs long simulations of this system, and found that JH2 and FERM came together to develop extensive interactions, giving rise to a compact four-domain model of JAK2 (Figure 1A, state 4) in which the FERM F3 subdomain is wedged between JH2 and JH1, and SH2 interacts with the C lobe of JH1. Importantly, F3 interacts with the αC helix and β4 strand of JH2, where the V617F residue is located (Figure 1A, state 4). In the model, V617 is in a hydrophobic cluster with the F285, W298, and L352 residues of F3, and the F595 and V615 residues of JH2 (Figure 1C). The model additionally features a pronounced electrostatic complementarity at the interface (∼3,500 Å^2^ buried surface area) between the JH2-JH1 and FERM-SH2 units (Figure 1D), which entails an array of salt bridges (Figure S1C). Using homology modeling, we constructed similar full-length models of JAK1, JAK3, and TYK2, and found that similar electrostatic complementarity exists for the other JAK proteins (Figures S1E), providing support for this full-length model of JAK2.

We subsequently added the SH2-JH2 linker to the model. Starting from a fully extended conformation, the linker settled into a collapsed conformation during our simulations, which did not use any artificial biasing force (Figure S1B, states 1–3). In this conformation, the phosphorylated S523 (pS523) residue of the linker is located at the center of the inactive model, in a highly positively charged region adjacent to the N-terminal end of the JH2 αC helix, where it interacts with the helix dipole and forms salt bridges with both JH2 (H587 and R588) and the FERM F3 subdomain (K302; Figure 1E, and Figure S1B, state 3). The energetically favorable interactions of pS523 suggest that, by mediating the JH2-JH1 and FERM-JH2 interactions, the phosphorylation of S523 “locks down” the four-domain structure; this may explain why phosphorylation of this residue is necessary to maintain a low basal activity of JAK2.^14^

When we simulated JH2 with the linker present but with S523 unphosphorylated, we found that S523 transiently approached the γ phosphate of the JH2-bound ATP (Figure S1D). This is consistent with the notion that JAK2 autoinhibition may occur with the protein first adopting the inactive conformation and cis-phosphorylation of S523 by JH2 subsequently stabilizing it.^14^ The functional importance of the SH2-JH2 linker^17^ position also goes beyond the pS523 position. In the linker conformation, the C-terminal region mostly packs with the FERM F3 subdomain; residues 501 to 505 (PPKPK) of the N-terminal region are between JH1 and JH2, mediating part of the JH2-JH1 interaction (Figure S1C). (In our previous model,^16^ the C-terminal region of the linker occupies the JH2 surface at its αC helix and β4 strand.) This is consistent with past findings that a cluster of mutations at the C-terminal region of the linker (e.g., M535I, H538L, K539L, an F537-K539 deletion, and H538Q/K539L) result in JAK2 activation and are associated with myeloproliferative neoplasms and other forms of cancer,^29–32^ and that some N-terminal mutations of the linker suppress the V617F phenotype.^33^

For completeness, we added the box1 and box2 regions of the Erythropoietin (EPO) receptor (EpoR), one of the key cytokine receptors with which JAK2 interacts, onto the JAK2 model based on homology modeling, using as templates the TYK2 and JAK1 crystal structures of FERM-SH2 bound with a segment of interferon-α receptor type 1 (IFNAR1)^11, 34^ (Figure S1B, state 4 and Figure 1A, state 5). Low-resolution electron microscopy (EM) envelopes^19^ have suggested that JAK1 is conformationally dynamic, transitioning between a highly compact conformation and an elongated one. We reasoned that this is likely to be the case for JAK2 also, and that the compact conformation probably entails tight packing between the FERM-SH2 and JH2-JH1 units that should reinforce the autoinhibition of JH1 by JH2 and thus is more likely to reflect the inactive state. It is thus reassuring that our full-length model of inactive JAK2 fits well into the EM envelope of the highly compact JAK1 conformation (Figure 1F).

### An active monomer conformation derived from simulations of the inactive monomer model

Release of JH2 and JH1 from the autoinhibitory structure was previously postulated to be integral to cytokine-induced JAK2 activation.^14, 16, 17^ Specifically, it was suggested that, upon activation, JAK2 dimerizes to facilitate auto-phosphorylation of both JH1s, which are released from their respective JH2s, while tethered by the long JH2-JH1 linker.^22^ It is likely that the release of JH1 is followed by a re-arrangement of the other three domains. In our inactive model, these three domains are broadly consistent with an EM envelope of JAK1 that is an intermediate between the compact and elongated envelopes^19^ (Figure S2C), and this EM envelope may thus reflect a state of JAK2 after JH1 release and prior to the rearrangement of the remaining three domains.

These considerations led us to simulate the three-domain construct of JAK2 starting from the inactive monomer conformation (with JH1 removed), and we introduced the V617F mutation based on the conjecture that the mutation helps to destabilize the inactive conformation^16^ (Figure 2A, state 1). In the simulation, we observed a conformational rearrangement in which JH2 pivoted on the FERM F3 subdomain and remained stable until the end of the simulation (Figure S2A). Upon this rearrangement, JH2 moved into the space that had been occupied by JH1 in the inactive model, and came into contact with SH2 while remaining in contact with F3 (Figure 2A, state 2). (In contrast, in the inactive model, JH2 interacts with F3 but not SH2, while JH1 interacts with both F3 and SH2.) With the repositioning of JH2, the SH2-JH2 linker settled into a conformation in which the N-terminal region of the linker docked to F3 and the C-terminal region of the linker was wedged between F3 and JH2 (Figure 2A, state 3, Figure 2B and Figure S2B). This linker conformation suggests that pathogenic exon 12 mutations in the C-terminal region of the linker may affect the F3-JH2 interaction and potentially stabilize the active conformation. We found this active conformation compelling, in part because it gave JAK2 an elongated shape consistent with the low-resolution EM data (Figure 2C). In simulations starting from this elongated conformation, we also observed a reversal of the rearrangement whereby JH2 returned from the putative active position to the inactive position (Figure S2A). This observation suggests a dynamic equilibrium between the active and inactive conformations for monomeric JAK2.

**Figure 2:**
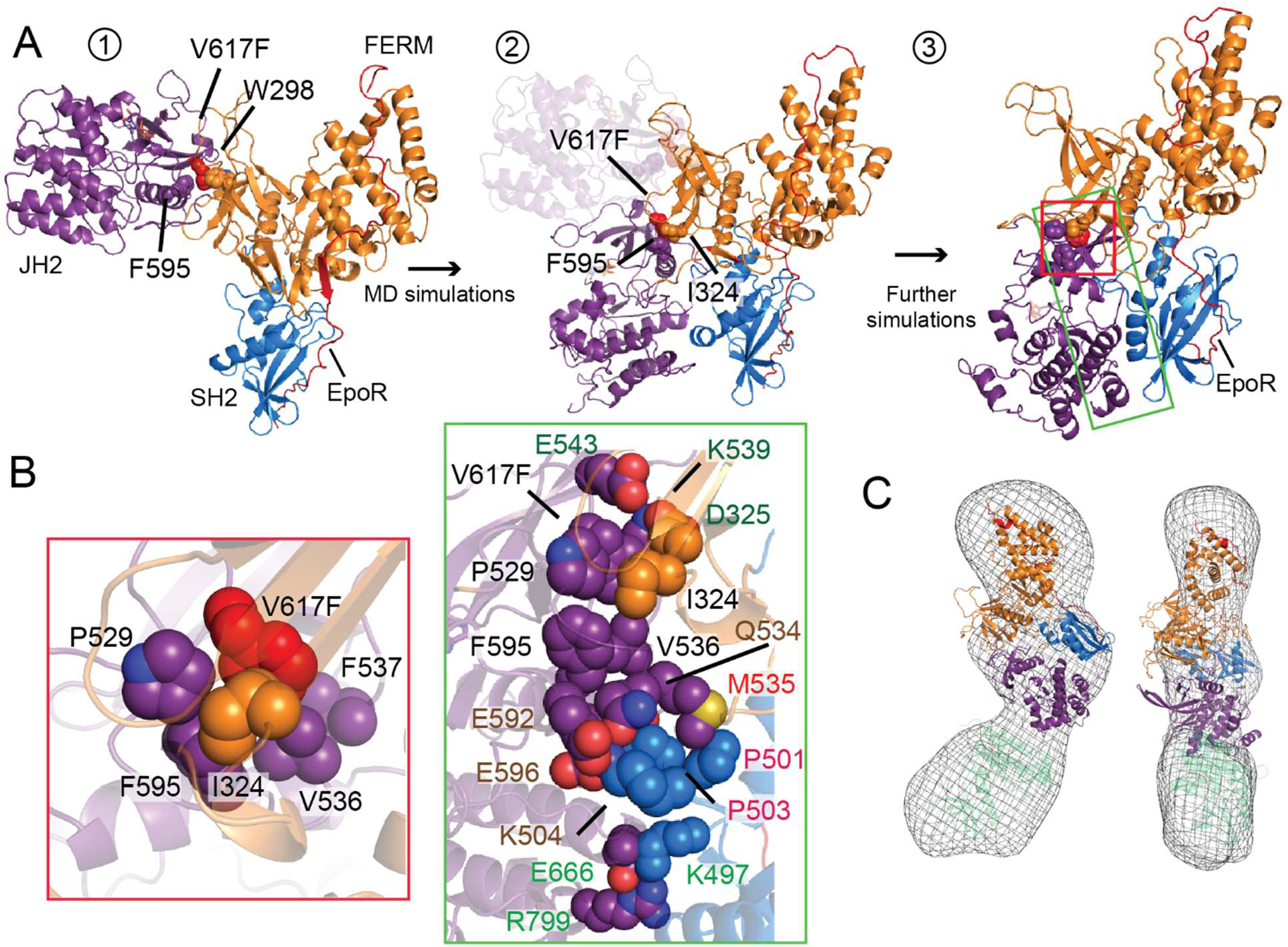
The active monomer model. (A) Generation of the active model: (1) the inactive monomer model with JH1 removed (and the V617F mutation introduced) as the simulation starting point; (2) repositioning of JH2 in a simulation; (3) the active monomer model from further simulations, with the V617F region and the JH2 interface with FERM-SH2 highlighted by rectangles. (B) Close-ups of the V617F region and the JH2 interface with FERM-SH2. (C) Fit of the active monomer model into a JAK1 EM envelope^19^ (front and side views), with JH1 arbitrarily placed to fill the envelope. Domains are colored as in Figure 1.

The F3 subdomain of FERM interacts closely with JH2 in both the active and inactive models, but the F3 interfaces with JH2 differ between the two. The V617F residue is centrally positioned at the F3-JH2 interface in both models. In the active model, the mutation residue belongs to a hydrophobic cluster composed of the P529, F595, and F617 residues of JH2 and the I324 residue of FERM (Figure 2B). It is likely that the V617F mutation reinforces this hydrophobic cluster, shifting the conformational equilibrium in favor of the active conformation.

### A model of the JAK2 active dimer mediated by JH2 and FERM

Upon ligand binding, cytokine receptors form dimers that lead to JAK2 activation.^35^ The constitutive activation of V617F requires dimerization of JAK2.^36^ We reasoned that ligand-induced dimerization of cytokine receptors may lead to dimerization of wild-type JAK2, and that the same active dimer structure may underpin the constitutive activation of V617F and the cytokine-induced activation of the wild-type JAK2. We thus aimed to assemble two JAK2 proteins into an active dimer configuration.

Ligand binding, receptor dimerization, receptor degradation, and JAK dimerization are closely coupled processes. In TYK2-deficient cells, the interferon receptor IFNAR1 loses its binding with interferon IFNα2,^37^ likely because TYK2-free IFNAR1 is incapable of dimerization. IFNα2 binding (which requires IFNAR1 dimerization) is not rescued by introducing a FERM-SH2 construct of TYK2 to the cells,^38^ suggesting that the absence of JH2 and JH1 precludes receptor dimerization. Furthermore, single-molecule imaging has shown that the absence of JAK1 reduces IFN-induced dimerization of IFNAR1 and IFNAR2.^39^ These past findings suggest that ligand-induced receptor dimerization is coupled to JAK dimerization, and that JAK dimerization is not entirely mediated by FERM and SH2. In view of JH1 needing to be set free for auto-trans-phosphorylation upon ligand binding and activation, JH2 is likely involved in mediating the dimerization of JAKs. This was previously suggested based on mutagenesis studies,^14, 22, 40, 41^ but the structural basis has remained unclear.

In seeking a structural model of the active dimer, we conducted 30 simulations, each 5 µs in length, starting from two spatially separated JH2 domains. These simulations generated a number of JH2 homodimer complex structures, but none of these structures led to a compelling JAK2 dimer. We then examined dimers present in reported crystal lattices of various FERM domains with the objective of identifying a FERM-FERM interface in an active JAK2 dimer. We constructed an active dimer model by superimposing our active monomer model on the crystal dimer of the FERM domain of focal adhesion kinase (FAK).^42^ Although the resulting JAK2 dimer model is indeed partially mediated by JH2 and allows simultaneous interaction with the membrane of the two FERM F2 subdomains, we eventually discarded this model because the dimerization interface appeared to be too small to sustain a stable dimer.

We then turned to mutation data on JAK2 in the Catalogue of Somatic Mutations in Cancer (COSMIC),^43^ a large compilation of mutations identified in cancer patients. We mapped the COSMIC JAK2 mutations onto our active and inactive monomer models, and found that more than half of the COSMIC mutations at the surfaces of the FERM, SH2, and JH2 units are located at the interdomain interfaces, or at the putative membrane interface at the F2 subdomain (Table S1). The other half of the COSMIC mutations are mostly located on the same side of the active monomer model (Figure 3A, red spheres), hinting at the possibility that these mutations occur at the active dimerization interface. An alternative explanation could be that these mutations affect the stability of the inactive JAK2 conformation, but this is unlikely according to the inactive monomer model, wherein they are solvent-exposed. Our active dimer model was thus informed by consideration of the COSMIC mutations, the FERM-membrane interaction, the assumed involvement of JH2, and the assumed symmetric nature of the dimer.

**Figure 3:**
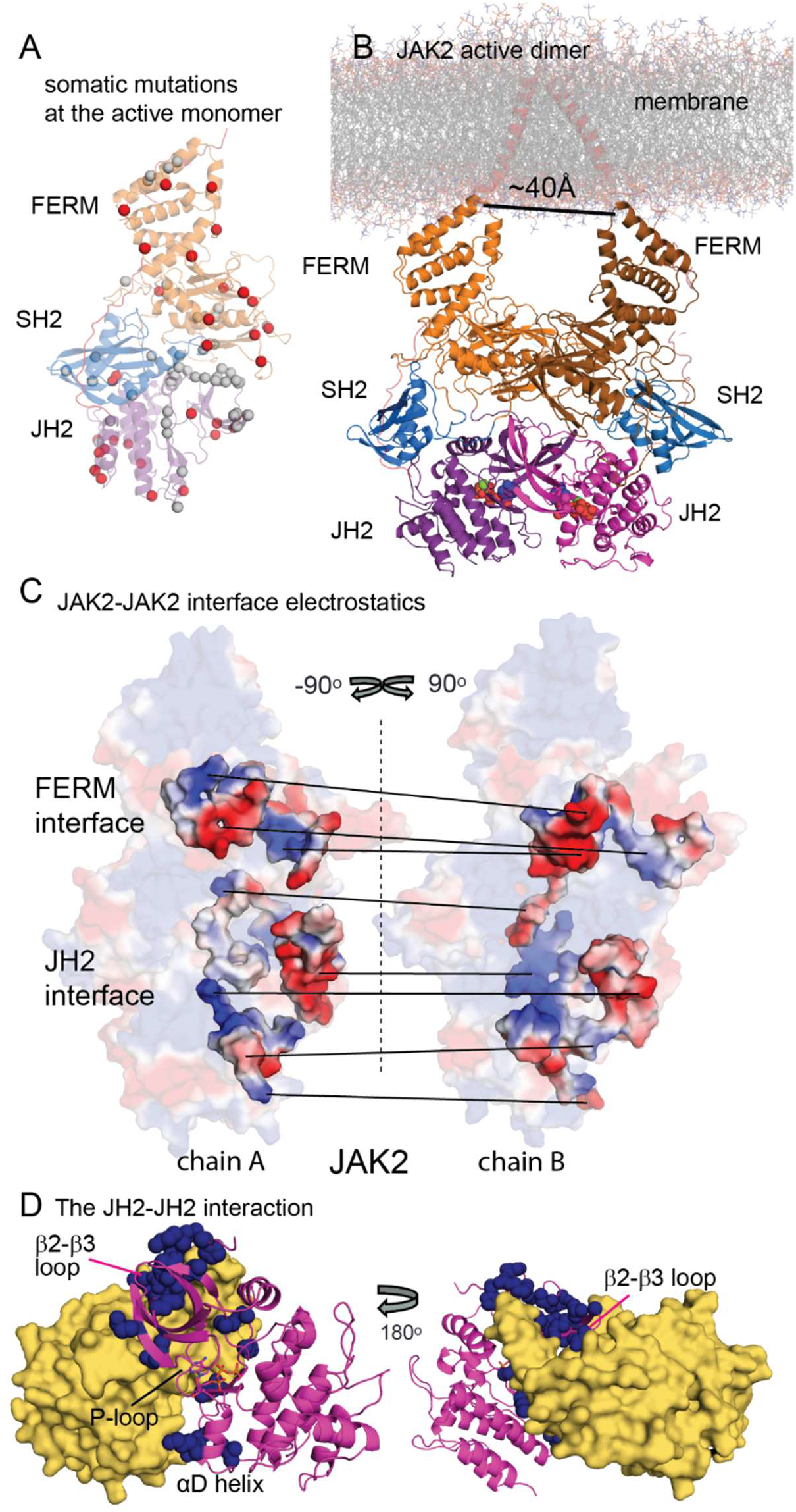
The active dimer model. (A) Mapping of somatic mutations onto the active monomer model. Mutations at the interfaces of inactive and active monomers or membrane interfaces are shown in gray, and other surface mutations are shown in red. (B) The active dimer model attached to a pair of EpoR peptides. JH2-bound ATPs are shown in space-filling representation. The separation of the C termini of the EpoR transmembrane helices is shown. (C) Surface electrostatics of the dimer interface. Straight lines connect the contacting regions at the interface. (D) The JH2-JH2 interaction in the active dimer. Somatic mutations at the JH2-JH2 interface are shown as dark-blue spheres. Except for panel D, domains are colored as in Figure 1.

We manually assembled two JAK2 active monomers to form a symmetric dimer model, in which one-third of unexplained COSMIC mutations are located at the dimerization interface. This dimer model was simulated for ∼11 µs, during which a well-packed dimer interface developed (Figure 3B). The dimer was highly stable, and we did not observe any large rearrangements in the simulation. In the dimer model after the simulation, the interaction between the two monomers is largely electrostatic in nature (Figure 3C), with a buried interface surface area of 4,900 Å^2^ (of which ∼2,100 Å^2^ is assigned to the JH2-JH2 interface and 2,300 Å^2^ to the FERM-FERM interface). By our design, the dimer interface is enriched with COSMIC mutations, with 14 located at the JH2-JH2 interface and 5 at the FERM-FERM interface (Figure 3D Figure S3A and Figure S3B). The FERM-FERM interface in our active dimer model is largely composed of loops that have relatively low sequence conservation among JAKs,^44^ and a few salt bridges (Figure S3C and S3D, Panel 1). This suggests that the FERM-FERM interaction may be weak and non-specific. In contrast to the FERM-FERM interface, the JH2-JH2 interface is well-packed, containing several salt bridges (Figure S3C and S3D, Panels 2-4).

Based on sequence alignment and our homology models of other JAKs based on the JAK2 dimer model, it is clear that the electrostatic features of the dimerization interface are mostly conserved (Figure 3C and Figure S3E), which is compatible with JAK heterodimerization. The JH2-JH2 interface primarily involves the “crossing” of the N-lobe β strands, especially the long β2-β3 sheets, at an angle of approximately 120° at the “backside” of JH2 (the side opposite to the ATP-binding site); it additionally involves the hinge region and αD helix (Figure 3D). The backside of a protein kinase domain is known to be dynamically coupled with the ATP-binding P-loop (i.e., the β1-β2 loop).^45^ This model suggests that ATP-binding by JH2 helps to maintain the high constituent activity of V617F^26^ by modulating the conformation of the backside.

A crystal structure of the extracellular module belonging to the ligand-induced active dimer of EpoR has shown that the extracellular C termini (the points of insertion into the membrane) are separated by 30–40 Å.^46–49^ Our active JAK2 dimer model predicts a separation of ∼40 Å between the N termini of the intracellular EpoR segments bound to the JAK2 dimer. This is geometrically highly compatible with the ligand-induced extracellular dimer structure of EpoR (Figure 3B). This active dimer model is also consistent with the observation in a fluorescence resonance energy transfer (FRET) study^50^ that extracellular ligand binding of growth hormone receptors modulates the proximity of the receptors’ juxtamembrane segments and the proximity of JAK2 FERM domains, giving rise to JAK2 activation. The geometry of our active dimer model is consistent with a C-terminal dimer model of the receptors’ transmembrane helices that was suggested in the study.

### A preliminary asymmetric inactive dimer model of JAK2

Pre-formed (ligand-free) EpoR dimers are observed experimentally under some cellular conditions^51, 52^ but not others.^21^ The ligand-free dimerization is likely to depend on the expression level and local EpoR density, and at a surface density of ∼0.3 molecule µm^−2^, pre-formed EpoR dimers are largely absent.^21^ A ligand-free dimer structure of the extracellular segment of EpoR has been captured crystallographically, in which the C termini are separated by a distance of 73 Å.^35^ It has been suggested that this crystal structure represents the structure of pre-formed EpoR dimers in cells at high EpoR density at the cell surface, and that EpoR ligand binding prompts JAK2 activation by reducing the C-termini separation to 30–40 Å.^35^ If so, two JAK2 proteins associated with a pre-formed EpoR dimer should be in an arrangement distinctively different from the active dimer to ensure low basal JAK2 activity. More recently, an activating surrogate ligand of EpoR was shown to lead to EpoR dimerization with a separation of more than 120 Å between the extracellular C termini.^21^ One interpretation is that JAK2 forms an autoinhibitory dimer at a C-terminal separation of ∼73 Å, whereas the 120-Å separation disrupts this autoinhibitory JAK2 dimer and increases JAK2 basal activity by allowing trans-phosphorylation of JH1, which is feasible given the length of JAK2.

Assuming the existence of an autoinhibitory JAK2 dimer, and building on the inactive monomer model of JAK2, we ventured to model the putative inactive dimer. Conjecturing that FERM and JH2 may interact in trans in this dimer, which would be consistent with the EpoR separation of 73 Å in the putative inactive EpoR dimer,^35^ we ran simulations of the association between FERM and JH2 (Figure S4A state 1) analogous to our simulations of SH2-JH1 association. We ran 20 independent, 10-μs unbiased MD simulations, each of which produced a model of the FERM-JH2 complex (Figure S4A, state 2). Based on the inactive monomer model of JAK2, each of the 20 FERM-JH2 models was extended into a full-length JAK2 dimer model. Models that lead to clashes between the two JAK2 inactive monomers were then eliminated, and focus was placed on a single inactive dimer model (Figure S4A, state 3 and 4). In this inactive JAK2 dimer model, the two EpoRs attached to the JAK2 dimer are separated by 90 Å at the C termini of the transmembrane helices. This separation is highly consistent with the ligand-free EpoR dimer structure,^35^ in which the C-terminal separation of the extracellular segments is 73 Å. In our inactive dimer model, the dimer interface is dominated by an interaction between FERM and the C lobe of JH2 in trans (2,500 Å^2^ buried surface area). FERM also interacts with the C lobe of JH1 in trans. This model allows for the possibility of two inactive JAK2 monomers both maintaining their interaction with the membrane through the F2 subdomain of FERM and C lobe of JH2. Ligand-independent oligomerization has been observed in EPO receptors,^52^ suggesting oligomerization of inactive JAK2. Intriguingly, our proposed inactive JAK2 dimer is asymmetric and can be extended into an oligomer while largely maintaining the orientation of each constituent monomer with respect to the membrane (Figure S4B).

### Somatic mutations are predominantly located at interdomain or dimerization interfaces in our models

The COSMIC database^43^ reports 633 somatic mutations of the JAK2 gene. These 633 somatic mutations include both point mutations and in-frame deletion or insertion mutations. Of the 633 somatic mutations, 448 are missense mutations that map to 291 mutation sites on JAK2, including 69 mutations mapped to the SH2-JH2 linker, six to V617, and 20 to either I682 or R683. Of these 291 mutation sites, we focused our analysis on the 107 surface mutation sites located on linkers or the surfaces of JH2, JH1, or the FERM-SH2 unit (Figure 4A, Table S1). Although these mutations are not necessarily gain-of-function and the mechanism of pathogenesis has yet to be established for many of them, we conjectured that many of these 107 mutations should be located at the cis or trans interdomain interfaces in either the active or inactive JAK2 structure, and that they should alter the conformational equilibrium and/or dimerization of JAK2. This notion is consistent with the fact that more than a quarter of these mutation sites (27 of the 107) are involved in the previously resolved JH2-JH1 interaction^16, 17^ (Figure 4A).

**Figure 4:**
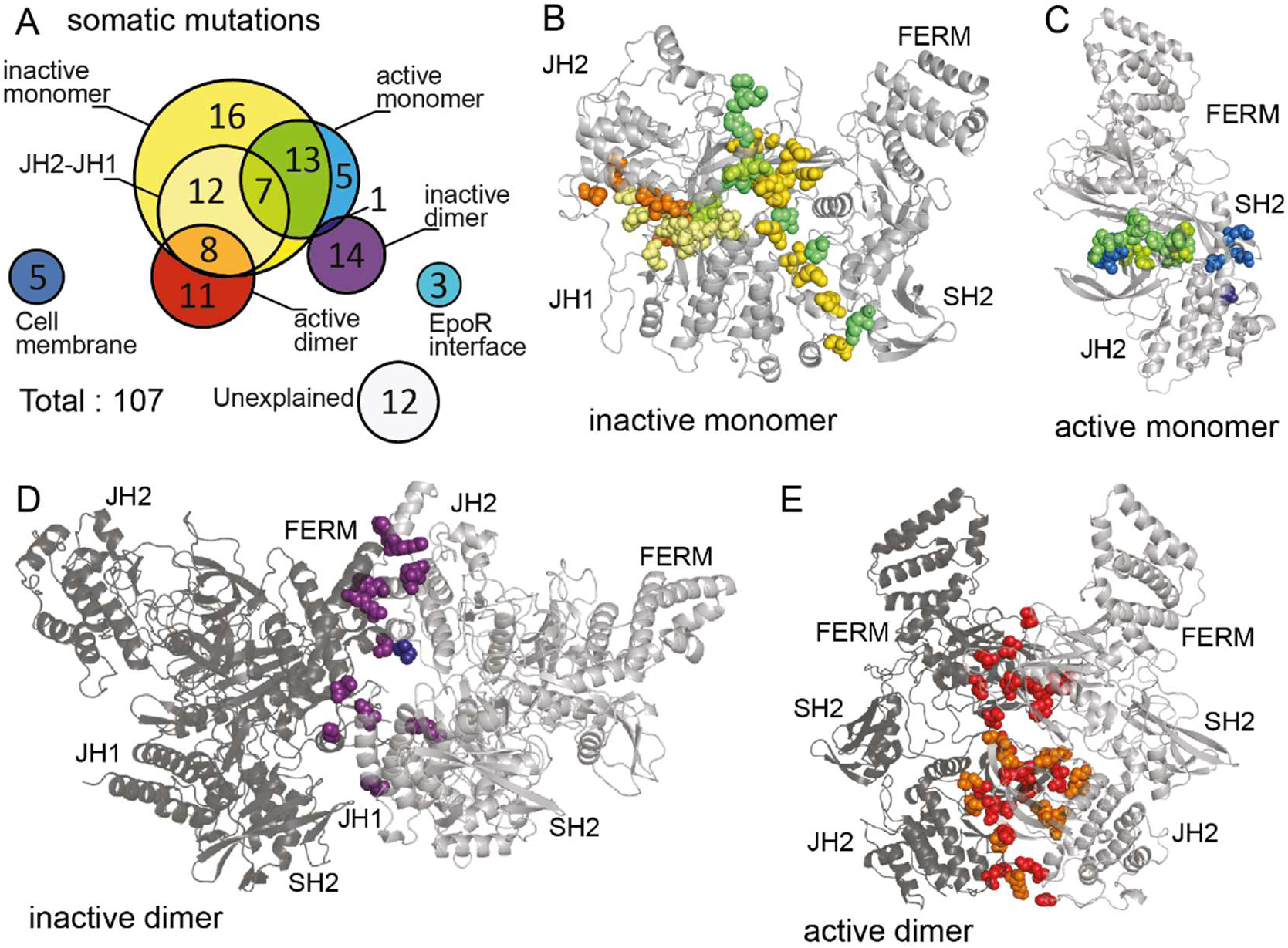
Somatic cancer mutations mapped to the models. (A) Venn diagram of somatic cancer mutations located at domain surfaces or at the SH2-JH2 linker. Each circle represents the mutations mapped to one or a set of interfaces and each number represents the counts of the mutations represented by an enclosed area. The locations of these mutations (see Table S1) can be seen by their coloring in the inactive monomer (B), the active monomer (C), the active dimer (D), and the inactive monomer (E) models.

We analyzed the COSMIC mutation sites in relation to our modeled interdomain interfaces (except for the FERM-SH2 interface, which is the same as in the crystal structures in all our models). Our analysis showed that 62 of the 107 surface mutation sites (58%) are located at the interfaces of the inactive and active monomer models (Figure 4B and Figure 4C), even though the interfaces together account for only 12% of the total surface area of JH2, JH1, and the FERM-SH2 unit. This represents a substantial enrichment of the COSMIC mutation sites at the interfaces. The inactive dimer model, which is more speculative, also represents a substantial enrichment at the dimer interface: It contributes another 13 surface COSMIC mutation sites (12%), despite the dimer interface accounting for only 5% of the total surface area (Figure 4E). These observations provide support for our inactive monomer, active monomer, and inactive dimer models, since these models’ construction was independent of the COSMIC data. We constructed the active dimer model partly based on the COSMIC data, and 19 COSMIC mutation sites are located at the dimer interface of that model (Figure 4D). Overall, the interdomain and dimerization interfaces of all four JAK2 models proposed here account for 87 of the total 107 surface COSMIC mutation sites (81%), despite these interfaces accounting for only 23% of the total surface area (Figure 4A, Table S2).

There are 56 mutation sites, including V617, at the cis interdomain interfaces in our inactive monomer model (Figure 4B), and roughly half (29) of them are clustered at the interface between the FERM-SH2 and JH2-JH1 units (Figure S5A). In the active monomer model, 25 mutation sites are located at the cis SH2-JH2 interface (Figure 4C). There are 20 mutation sites that belong to the interdomain interfaces of both the active and inactive monomer models; this is not surprising, given the significant overlap in the interfaces of the inactive and active monomers. A smaller number of mutation sites (15) are located at the dimerization interface of the inactive dimer model (Figure 4D).

Of the 20 surface mutation sites that are not located at any of the interdomain or dimerization interfaces in our structural models, 8 appear likely to interfere with the receptor binding or membrane interaction of JAK2. Four of these 8 mutation sites are arginine residues at the α3 helix of FERM, and these mutations may disrupt membrane localization of JAK2;^53^ R761 is another such mutation site at the αG helix of JH2, which may interact with the membrane according to our inactive monomer model (Figure S5B). The other 3 of these 8 mutation sites are located at a four-strand β sheet that is involved in interactions between JAK2 and cytokine receptors^11^ (Figure S5B, Table S1). (It remains unclear how these mutations are related to cancer growth, since they appear likely to abolish the activity of JAK2 by disrupting its membrane localization or receptor binding.)

### Mutagenesis experiments support the JAK2 models

Our models predicted a FERM-JH2 interface and an SH2-JH1 interface in the inactive monomer of JAK2, and another FERM-JH2 interface in the active monomer. Consistent with this prediction, our mutagenesis experiments showed that a number of JAK2 mutations at these interfaces (Figure 5A) modulated the constitutive activity of V617F. Unlike previously identified V617F-rescuing mutations^23, 24, 33, 41^ located at the N lobe of JH2 or at the SH2-JH2 linker, the mutations we identified here are novel in that they are located on the FERM unit or C lobe of JH2. The mutagenesis experiments provide compelling evidence for our models, in which FERM plays an important structural role in both the active and inactive conformations.

**Figure 5:**
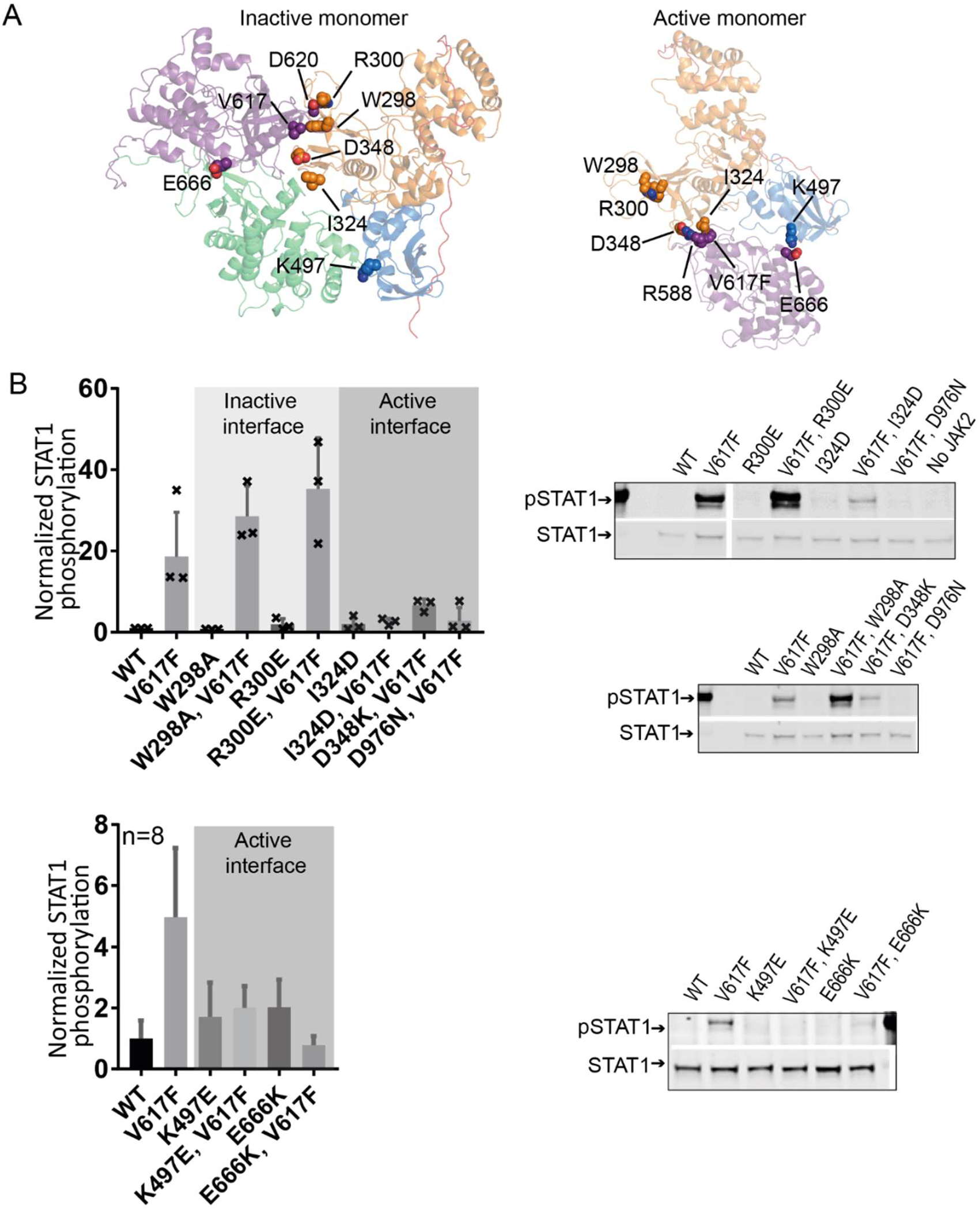
Mutagenesis experiments on residues at the inactive and active monomer interfaces. (A) The mutated residues mapped to the interfaces of the monomer models. (B) At left, pSTAT1 phosphorylation (normalized by STAT1 level) of JAK2 mutants as fold change relative to basal phosphorylation of JAK2 wild-type: The upper left shows average values and standard deviations from three independent experiments, with each individual value denoted by an x; the lower left shows average values and standard deviations from eight independent experiments (individual values are not marked because of the larger data set). At right, representative western blots of immunoprecipitated JAK2 from ϒ2A cells. Original images of blots used in this study can be found in Figures S6 and S7.

The V/F617 residue in our active monomer model is part of a hydrophobic cluster that comprises the P529, F595, and V/F617 residues of JH2 and the I324 residue of the FERM F3 subdomain (Figure 2B). From this observation and the high constitutive activity of V617F, we inferred that the V617F mutation stabilizes this hydrophobic cluster, and the F595A/V mutations, which rescue the V617F mutant,^23, 24^ may counter the stabilizing effect. We tested the I324D mutation against the background of the V617F mutant and found that, as expected from the active model, I324D significantly suppressed the constitutive activity of the V617F mutant in terms of phosphorylation of both the JH1 activation loop (pTyr1007 and pTyr1008) and downstream STAT1 (Figure S6A and S6B). Surrounding the hydrophobic cluster in the F3-JH2 interface are several salt bridges between F3 and JH2 (Figure 2B), including E596-K504 and D348-R588. The E596-K504 salt bridge is consistent with the E596A and E596R mutations suppressing V671F activity.^33^ Our experiments showed that D348K indeed also suppresses the constitutive activity of V617F (Figure 5B). In the active monomer model, E666 (of JH2) forms a salt-bridge with the K497 residue (of SH2) at the SH2-JH2 interface; our experiments showed that E666K and K497E mutations each suppressed the constitutive activity of V617F (Figure 5B; Figure S6A and S6B; Figure S7) to a level similar to the kinase-dead D976N mutation. ^41^ The I324, D348, K497, and E666 residues are largely exposed in the inactive model and do not appear to be involved in important and stable inter-residue interactions, and they are unlikely to affect the stability of the inactive conformation. Together, these experimental findings provide strong support for our active monomer model.

In the inactive monomer model, V/F617 is located at the interface of JH2 and F3, where V/F617 and the F595 residue of JH2 share a hydrophobic cluster with the F285, W298, and L352 residues of the FERM F3 subdomain (Figure 1C and Figure S1C). To validate the inactive monomer model, we tested the W298A mutation experimentally. W298A is solvent-exposed in the active models and unlikely to directly stabilize the active structures of JAK2. We found that, as predicted by the inactive model, the W298A mutation further raised the high constitutive activity of the V617F mutant (Figure 5B; Figure S6A and S6B), likely by impairing the packing of the hydrophobic cluster and further destabilizing the inactive conformation. In addition to the hydrophobic packing, in the inactive monomer model, JH2 also interacts with the F3 subdomain through an array of salt bridges (Figure S1C), including one between R300 and D620. This is supported by our experimental finding that R300E (which is solvent-exposed in the active dimer) also raised the constitutive activity of V617F (Figure 5B). Our experiments showed that W298A and R300E individually were not sufficient to raise the basal activity of wild-type JAK2, suggesting that (for the wild-type) destabilizing the inactive conformation without stabilizing the active conformation may not lead to higher activity.

## Discussion

Starting from the two resolved two-domain structures of JAK2—the JH2-JH1 structure^16, 17^ and the FERM-SH2 structure^11^—we constructed full-length models of JAK2, including an inactive monomer model, an active monomer model, and an active dimer model (Figure 6A). In view of evidence that EpoR dimerizes without ligand binding and JAK2 proteins form inactive dimers under certain cell conditions,^51, 52^ we also extended the inactive monomer model to a more speculative inactive dimer model (Figure 6B). Low-resolution negative-stain EM images of JAK1 have captured a series of conformations from highly compact (“closed”) to elongated (“open”).^19^ Our inactive and active JAK2 monomer models are consistent with the closed and open envelopes, respectively (Figure 1F and Figure 2C). In the inactive model, JH1 is sequestered from auto-phosphorylation, whereas in the active model, JH1 is unconstrained and tethered to JH2 with a long linker. The EM envelopes and our simulations both suggest that JAK2 exists in dynamic equilibrium between closed and open conformations. The models provide insights into both JAK2 activation and autoinhibition and the constitutive activity of the V617F mutant.

**Figure 6:**
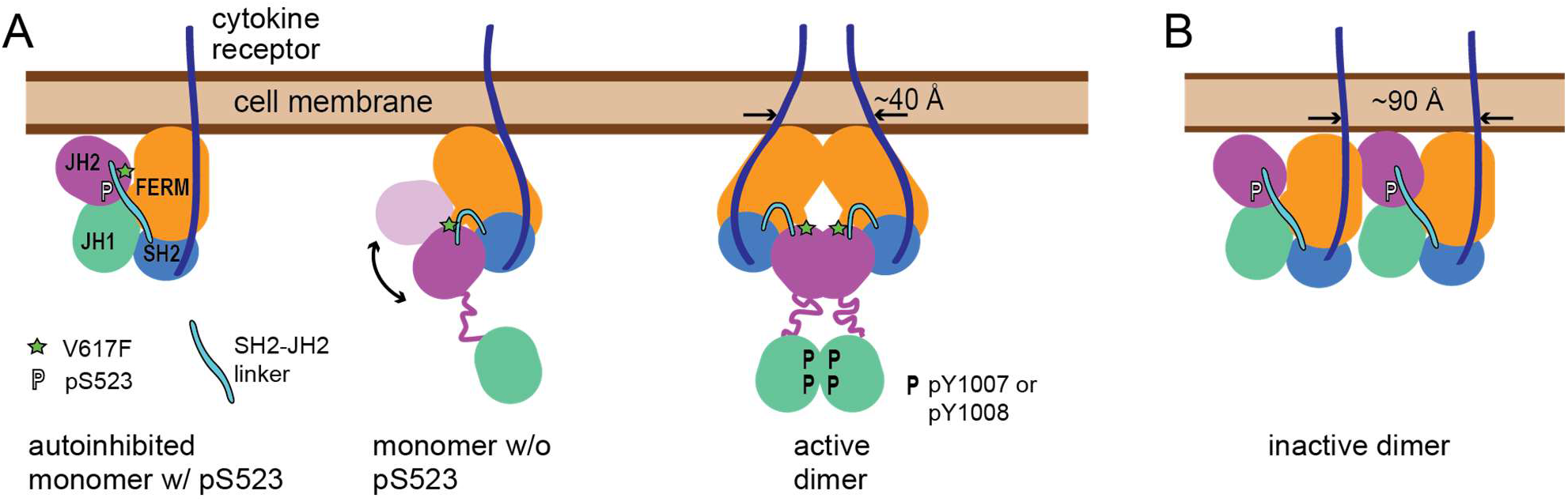
Cartoon summary of the structural models. (A) The three main states of JAK2: the monomer with phosphorylated S523 (pS523) locked in the inactive conformation, the unphosphorylated monomer transitioning between the active and inactive conformations, and the active dimer. The separation of the C termini of the transmembrane helices of the two cytokine receptors is labeled. Phosphorylated residues are denoted by the letter “P”. (B) The more speculative inactive JAK2 dimer model.

Our model of the active dimer suggests how cytokine receptors and JAK2 are conformationally coupled: The separation of the C termini of the transmembrane helices in a ligand-bound EpoR dimer is ∼40 Å, which is highly compatible with the separation of 30–40 Å between the C termini of the extracellular segments in the EpoR dimer.^46–49^ In contrast, in our inactive dimer model, the separation between the C termini of the EpoR transmembrane helices is much larger (∼90 Å). Recently, based on a crystal dimer of the FERM-SH2 units of JAK2 bound to an EpoR peptide, an intriguing active dimer model was proposed,^54^ but this model suggests a separation of 120 Å or more between the receptor transmembrane helices, and so it is not clear whether this model is compatible with the ligand-bound EpoR dimer structure.

Our inactive monomer model suggests a mechanism for phosphorylation-dependent JAK2 autoinhibition. In the model, phosphorylated S523 is centrally positioned in the model and “locks down” the inactive conformation together with the SH2-JH2 linker (Figure 6A). pS523 is located near the N terminus of the JH2 αC helix, and the linker is positioned toward the C lobe, wedged into a poorly packed region of JH2 between the αC helix and the activation loop of JH2.^25, 40^

Since the prevalent pathogenic mutation V617F is located at the center of the FERM-JH2 interface in both our active and inactive monomer models, we suggest that the unusually high constitutive activity of the V617F mutant may arise from the V617F mutation both weakening the inactive conformation and stabilizing the active conformation of JAK2. This is consistent with a recent analysis of mutations that suppress the activity of V617F.^41^ In our active dimer model, the F617 residue is packed with F595 and F537, and E592 forms a salt bridge with K415 (of SH2), while E596 forms a salt bridge with K502 (of SH2). These observations offer a structural explanation for the rescuing effect of the F595A, F537A, E592R, and E596R mutations.^23, 24, 33, 41^ I682 and R683 are two other sites of prevalent pathogenic mutations. Similarly to V617, these two JH2 residues are located at the inter-domain interface in our model of the inactive dimer, interacting with the β2-β3 loop of JH1. Additionally, I682 and R683 are located at the JH2-JH2 interface in our active dimer model, where those on one JH2 interact with the hinge region of the other JH2. Our models thus suggest that, similarly to V617F, the pathogenicity of the I682 and R683 mutations may arise from their dual effect on the stability of the active and inactive conformations of JAK2.

In addition to maintaining JAK2 autoinhibition,^13–15^ the JH2 pseudokinase domain also plays an important role in enabling full JAK activation.^55–57^ The models of JAK2 we have presented here provide a structural interpretation for this finding, by showing a central role for JH2—and, in particular, its long β2-β3 strands that are key to JH2-JH2 interactions—in the active dimer. Given the coupling of the β2-β3 sheets with the P loop of the ATP binding site,^45^ the active dimer model also offers a potential explanation for the observation that ATP binding is necessary for maintenance of the high constitutive activity of V617F. Our models suggest, moreover, that FERM is integral to JAK2 regulation. In the inactive monomer model, the F3 subdomain is wedged between JH2 and JH1, whereas in the active monomer model, F3 interacts extensively with JH2. Our identification of novel FERM mutations that alter the constituent activity of V617F lends strong experimental support to our structural models.

Our JAK2 models have significant implications for drug discovery. They suggest, for example, that the β2 and β3 sheets of JH2 are central to both the JH2-JH1 interaction in the inactive conformation and the JH2-JH2 interaction in the active dimer. The P loop of the ATP-binding site and the β2-β3 sheet likely form a conformational “seesaw” that pivots around the hinge (Figure S5C). By this mechanism, ATP binding of JH2, which directly affects the conformation of the P loop, would likely indirectly modulate the conformation of the β2-β3 sheet. In view of the suppression of V617F activity associated with disruption of ATP binding by JH2, the apo state of JH2 may favor the inactive interaction and disfavor the active interaction. We thus suggest that a small-molecule drug targeting the ATP-binding site of JH2 could counter the effect of the V617F mutation by reinforcing the apo-like conformation of the P loop. Additionally, our inactive monomer model suggests that JAK2 drug discovery programs could target grooves of the interface between the FERM-SH2 and JH2-JH1 units in the inactive monomer with the aim of stabilizing the inactive conformation of V617F.

## Methods

### Molecular dynamics simulations

All simulations were based on X-ray structures of the JAK2 FERM-SH2 unit (PDB code: 4Z32),^58^ the JH1 domain (PDB code: 3KRR),^59^ and our previously published model of the JH2-JH1 complex.^16^ The EpoR residues were built by homology modeling based on crystal structures of the FERM-SH2 unit in TYK2 and JAK1, each bound to an IFNAR1 segment.^11, 34^ Non-protein atoms and protein tags were removed, and with the exception of the SH2-JH2 linker, the missing loop regions and sidechain atoms were modeled to make a complete protein structure using the software package Maestro (Schrödinger, LLC). Simulation systems were set up by placing various systems in a cubic simulation box (with periodic boundary conditions) with at least a 10-Å distance from the protein surface to the edge of the simulation box. Na^+^ and Cl^-^ ions were added to obtain a neutral total charge for the system and maintain physiological salinity (150 mM). Explicitly represented water molecules were added to fill the simulation box. The systems were parameterized using the CHARMM36 force field^60, 61^ and the TIP3P water model,^62^ then equilibrated in the NPT-NVT scheduled ensemble at 1 bar and 310 K. Production MD simulations were performed on the special-purpose supercomputer Anton 2^63^ in the NVT ensemble at 310 K using the Nose-Hoover thermostat^64^ with a time step of 2 fs. All bond lengths to hydrogen atoms were constrained using an implementation^65^ of M-SHAKE.^66^ The Lennard-Jones and Coulomb interactions in the simulations were calculated using a force-shifted cutoff of 12 Å.^67^ The simulation trajectories were visualized and analyzed using Visual Molecular Dynamics (VMD) software^68^ and the images of protein structures were made using the PyMOL Molecular Graphics System (Schrödinger, LLC). All models went through a final restrained energy minimization using Maestro. The simulations reported here are described in Table S3.

### Transfection and western blot analysis

JAK2-deficient ϒ2A human fibrosarcoma cells were transfected with different human JAK2-hemagglutinin (HA) constructs in pCIneo vector (100 ng per 12-well plate well) with FuGENE HD (Promega). After 48 hours, cells were washed with ice-cold phosphate buffered saline (PBS), lysed in cold Triton X-100 lysis buffer with protease and phosphatase inhibitors (2 mM vanadate, 1 mM phenylmethanesulfonyl fluoride, 8.3 µg ml^−1^ aprotinin, and 4.2 µg ml^−1^ pepstatin), spun for 20 minutes at 16,000 g at 4°C, and the resulting supernatants were run on Mini-PROTEAN® TGX™ Precast Gels (BioRad). Immunoblots were blocked with bovine serum albumin (BSA) and incubated with primary antibodies for HA Tag (1:2000, OAEA00009, Aviva Systems Biology), phosphorylated JAK2 (pJAK2; 1:1000, 07-606, MilliporeSigma), phosphorylated STAT1 (pSTAT1; 1:1000, 7649, Cell Signaling), STAT1 (1:1000, 610116, BD Biosciences), or actin (1:1000, MAB1501R, MilliporeSigma) and a mixture of goat anti-rabbit and goat anti-mouse DyLight secondary antibodies (both from Thermo Fisher Scientific). Blots were scanned with an Odyssey CLx (LI-COR Biosciences), and immunoblot signals were quantified with Image Studio software (LI-COR Biosciences) by manually assigning bands and dividing the phosphorylation (pJAK2 or pSTAT1) signal values with the expression (HA or STAT1) signals.

## Data availability

The molecular dynamics trajectories described in this work are available for non-commercial use through contacting. Atomic coordinates from the reported models are also available upon request.

## Acknowledgments

The authors thank Michael Eastwood for helpful discussions and Jessica McGillen for editorial assistance. H.M.H., J.R., and O.S. were supported by the Academy of Finland, Sigrid Jusélius Foundation, Finnish Cancer Foundation, Jane and Aatos Erkko Foundation, Tampere Tuberculosis Foundation, and Pirkanmaa hospital district competitive research funding; S.R.H. was supported by National Institutes of Health grant R01AI101256.

