## Supplemental Information for "Structural models of full-length JAK2 kinase"

### Supplementary Tables

| intradomain interfaces | | | | | | | |
|---|---|---|---|---|---|---|---|
| JH1-JH2 | inactive monomer model | active monomer model | inactive dimer model | active dimer model | | not on interface | |
| N533 | E274 | I324 | G48 | V63 | FERM-SH2 | R130 | EpoR interactions |
| M535 | G281 | K414 | D53 | T108 | | H134 | |
| F537 | L316 | Q419 | D194 | N337 | | T514 | |
| H538 | Y317 | E442 | S203 | R340 | | K167 | |
| E566 | D319 | K497 | R215 | E386 | | E271 | |
| V567 | I324 | P503 | P727 | N548 | | L314 | |
| D569 | K346 | P529 | K728 | S550 | | R228 | cell membrane interactions |
| G571 | D348 | N533 | Q764 | R564 | | R230 | |
| E592 | K497 | M535 | R769 | E566 | | R234 | |
| S593 | P503 | F537 | H770 | V567 | | R235 | |
| S605 | P529 | H538 | A778 | D569 | JH2 | R761 | |
| H606 | K539 | K539 | N782 | G571 | | F560 | |
| K607 | R541 | I540 | S797 | S605 | | W777 | |
| E666 | N542 | R541 | K912 | K607 | JH1 | S887 | |
| R683 | E543 | E543 | T969 | L611 | | E889 | |
| K688 | F547 | H587 | | K640 | | K914 | |
| E846 | H587 | E592 | | R683 | | R923 | |
| L855 | E592 | S593 | | K688 | | L925 | |
| G858 | S593 | F595 | | A756 | | G1093 | |
| R867 | F595 | E596 | | | | S1115 | |
| P870 | E596 | C616 | | | | | |
| D873 | V617 | V617 | | | | | |
| T875 | C618 | C618 | | | | | |
| V878 | E621 | E621 | | | | | |
| Y931 | K945 | E666 | | | | | |
| P933 | E1052 | S797 | | | | | |
| N986 | K1055 | | | | | | |
| | A1059 | | | | | | |
| | R1063 | | | | | | |
| | D1068 | | | | | | |
| | G1086 | | | | | | |

**Table S1: List of JAK2 somatic cancer mutations.** Residues shown in Figure 4, listed and denoted using the same color scheme.

| | Inactive | | | | |
|---|---|---|---|---|---|
| | | ASA complex (Å²) | BSA (Å²) | ASA 1 (Å²) | ASA 2 (Å²) |
| Monomer | *JH2/JH1* | 32334 | 3414.8 | 17836 | 17913 |
| | *FERM-SH2/JH2-JH1* | 55565 | 3514.1 | 26744 | 32334 |
| Dimer | *JAK2/JAK2(full length)* | 110006 | 2539.4 | 56608 | 55937 |
| | Active | | | | |
| Monomer | *FERM-SH2/JH2* | 41301 | 2068.8 | 27565 | 15805 |
| Dimer | *JAK2/JAK2(3-domain)* | 78045 | 4921.6 | 41666 | 41300 |

**Table S2: Buried surface area (BSA)and accessible surface area (ASA) calculations for the models.** Shown are the accessible surface area ("ASA complex") and buried surface area of the given complex, and the accessible surface areas of the individual components in the unbound form.

| Simulation length | Box size | Number of atoms | Referenced figure | System description | Motivation | Restraints |
|---|---|---|---|---|---|---|
| 30 X 5 μs | ~95 Å | ~80K | Figure S1F | JH2 residues 536-809 and SH2 397-500 and ATP | JH2-SH2 unbiassed association | |
| 20 X 3 μs | ~100 Å | ~95K | Figure 1A, 1 | JH1 residues 839-1131 and SH2 397-500 | JH1-SH2 unbiassed association | |
| 8 X 10 μs | ~145 Å | ~290K | Figure 1A, 3 | FERM-SH2 residues 37-500 and JH1-JH2 residues 539-1131 (pT570) | Generation of 4-domain JAK2 inactive model | |
| 10 X 3 μs | ~118 Å | ~170K | Figure 1A, 4 | FERM-SH2 residues 37-500 and JH1-JH2 residues 523-1131 (pS523,pT570) | Modeling of of SH2-JH2 linker residues pS523-538 | |
| 10 X 3 μs | ~120 Å | ~175K | Figure 1A, 4 | FERM-SH2 residues 37-500 and JH1-JH2 residues 523-1131 (pS523,pT570) | Modeling of of SH2-JH2 linker residues 522-501 | |
| ~600μs | ~120 Å | ~180K | Figure 1A, 5 | JAK2 residues 37-1131 (pS523,pT570) and EpoR residues 273-293, 318-336 | Refinement of the full length inactive monomer model after modelling of EpoR | on EpoR* |
| ~30μs | ~120 Å | ~180K | | JAK2 residues 37-1131 (pS523,pT570) and EpoR residues 273-293, 318-336 and ATP | Refinement of the full length inactive monomer model after modelling of ATP | on EpoR and ATP* |
| 3 X 10μs | ~75 Å | ~45K | Figure S1D | JH2 residues 518-810 (S523) and ATP | S523 can access to JH1 ATP Υ-phosphate spontaneously for phosphorilation in cis | on ATP* |
| 20 X 10 μs | ~110 Å | ~130K | Figure S2A, 1 | FERM residues 37-405 and JH2 residues 536-809 | FERM-JH2 unbiassed association | |
| 10 X 2 μs | ~215 Å | ~995K | Figure S2A, 3 | 2 X JAK2 residues 37-1131 (pS523,pT570) and EpoR residues 273-293, 318-336 connected by 6-Gly linker | Generation of the JAK2-JAK2 inactive dimer model | on EpoR* |
| ~24 μs | ~215 Å | ~995K | Figure S2A, 4 | 2 X JAK2 residues 37-1131 (pS523,pT570) and EpoR residues 273-293, 318-336 connected by 6-Gly linker | Refinement of the JAK2-JAK2 inactive dimer model | on EpoR* |
| ~200 μs | ~118 Å | ~170K | Figure2A, 1 | FERM-SH2-JH2 residues 38-819 (S523, T570, V617F), EpoR residues 273-293, 318-336 connected by 6-Gly linker and ATP | Generation of active monomer model | on EpoR and ATP* |
| ~300 μs | ~122 Å | ~185K | Figure S3A | FERM-SH2-JH2 residues 38-819 (S523, T570, V617F), EpoR residues 273-293, 318-336 connected by 6-Gly linker and ATP | Spontaneous regeneration of the inactive monomer interface | on EpoR and ATP* |
| 11 μs | ~150 Å | ~340K | Figure 3B | 2 X FERM-SH2-JH2 residues 38-819 (S523, T570, V617F), EpoR residues 273-293, 318-336 connected by 6-Gly linker and ATP*** | Refinement of active dimer model | on EpoR, ATP*, domain-domain interfaces ** |

*distance restraints to keep the given molecule stable in simulation

**distance restraints to keep interfaces intact in the beginning of the simulation, these restraints were later released

***the MD simulations were run without lipids

**Table S3: Description of MD simulation systems reported.** Details of various MD simulations reported here and in the main text.

### Supplementary Figures

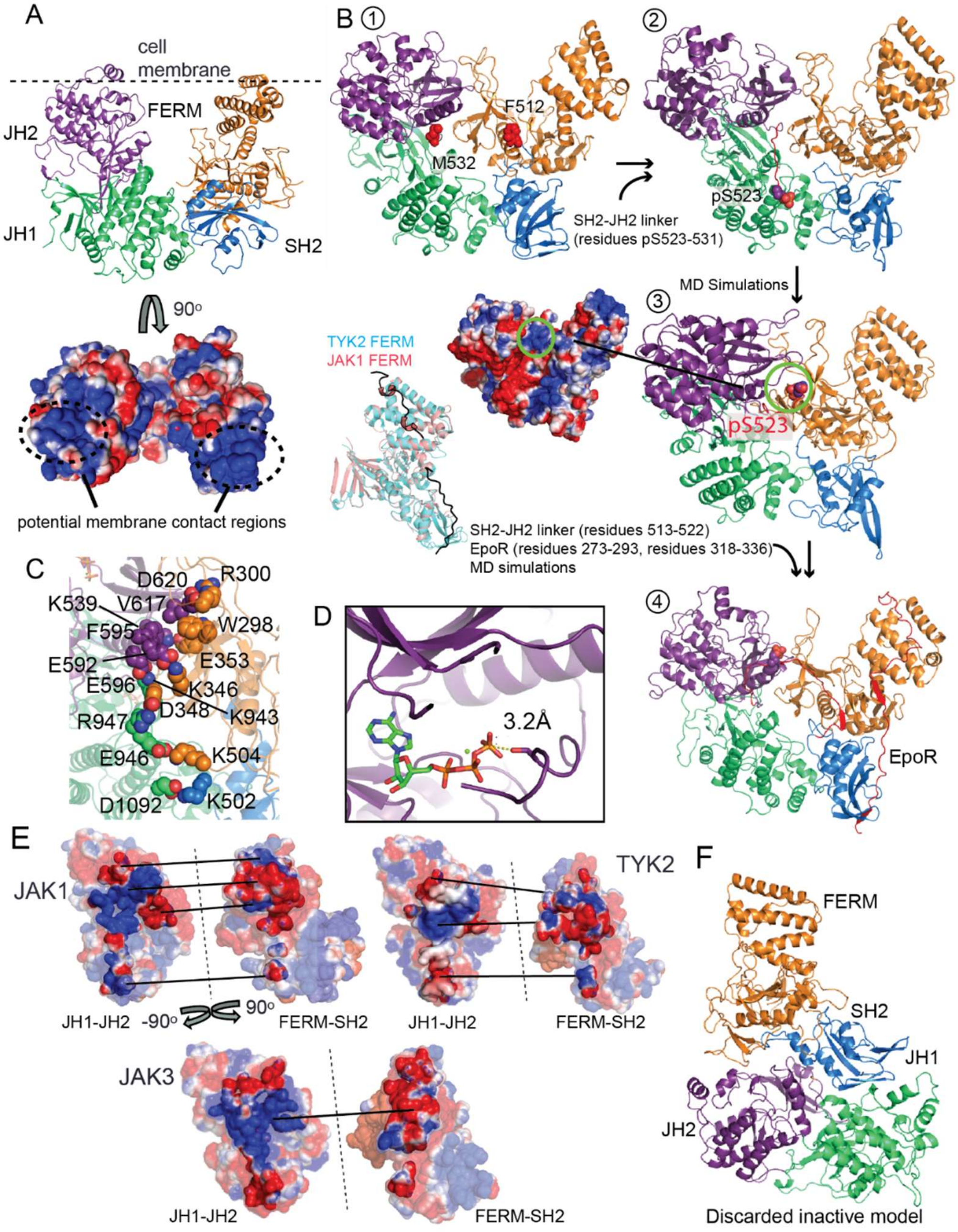


**Figure S1. Related to Figure 1: The inactive monomer model.** (A) (top) Initial inactive model (Figure 1A, step 3) and (bottom) its electrostatic features at the putative membrane binding interface. (B) Modeling of the SH2-JH2 linker and the EpoR receptor: (1) N-and C-terminal residues of the linker (red) in the inactive model (Figure 1A, step 4); (2) adding the C-terminal segment of the linker (residues pS523–531(red); pS523 in sphere representation) in an extended conformation; (3) the location of pS523 and electrostatics of the region (4) simulation-generated conformation after addition of the N-terminal segment of the linker (residues 513–522) and the EpoR peptide; the side panel (to the left of step 3) shows the crystal structure of the FERM-SH2 unit of TYK2 and JAK2 bound to cytokine receptor peptides, which are templates for our model of the EpoR peptide. (C) Close-up of the FERM-SH2/JH2-JH1 interface in the inactive monomer model. (D) The ATP and S523 residue of JH2 are transiently adjacent in simulations, potentially allowing cis phosphorylation of S523. (E) Electrostatic properties of the interface between the FERM-SH2 and JH2-JH1 units in the inactive models of JAK1, TYK2, and JAK3, which we derived from the JAK2 model by homology modeling. Each straight line denotes two matching regions at an interface. (F) An alternative model of the JAK2 inactive monomer that we eventually discarded. In all panels, domains are colored as in Figure 1.

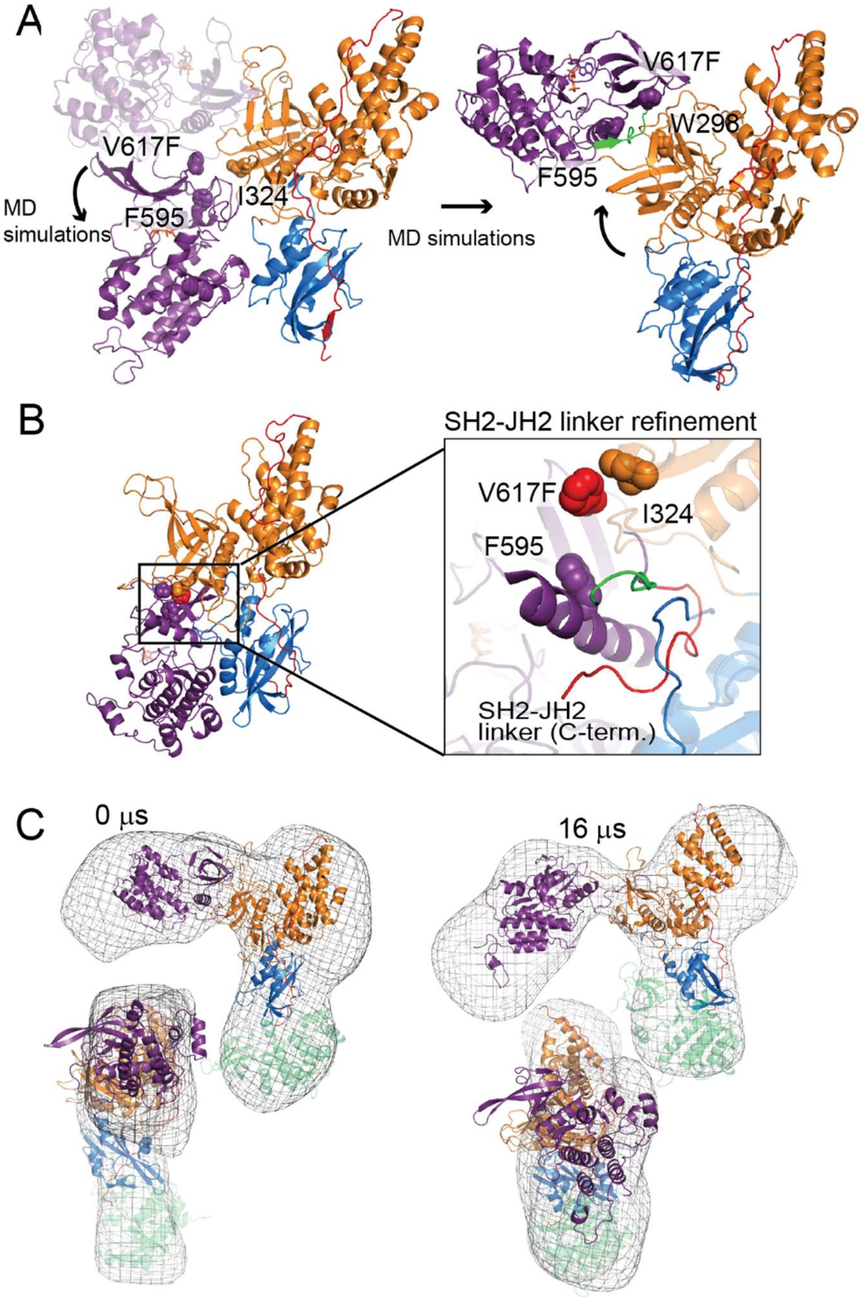


**Figure S2. Related to Figure 2: The active monomer model.** (A) JH2 returned to the inactive position in simulations of FERM-SH2-JH2 starting from the active conformation. ATP is shown in stick representation. (B) A close-up view of the FERM-JH2 interface and the SH2-

JH2 linker conformation in the active monomer model. The starting (red) and final (green) positions of the SH2-JH2 linker are shown in cartoon representation. (C) Conformations generated by the FERM-SH2-JH2 simulation fitted into EM envelopes of JAK2 (front and side views). JH1 is tentatively placed. Domains are colored as in Figure 1.

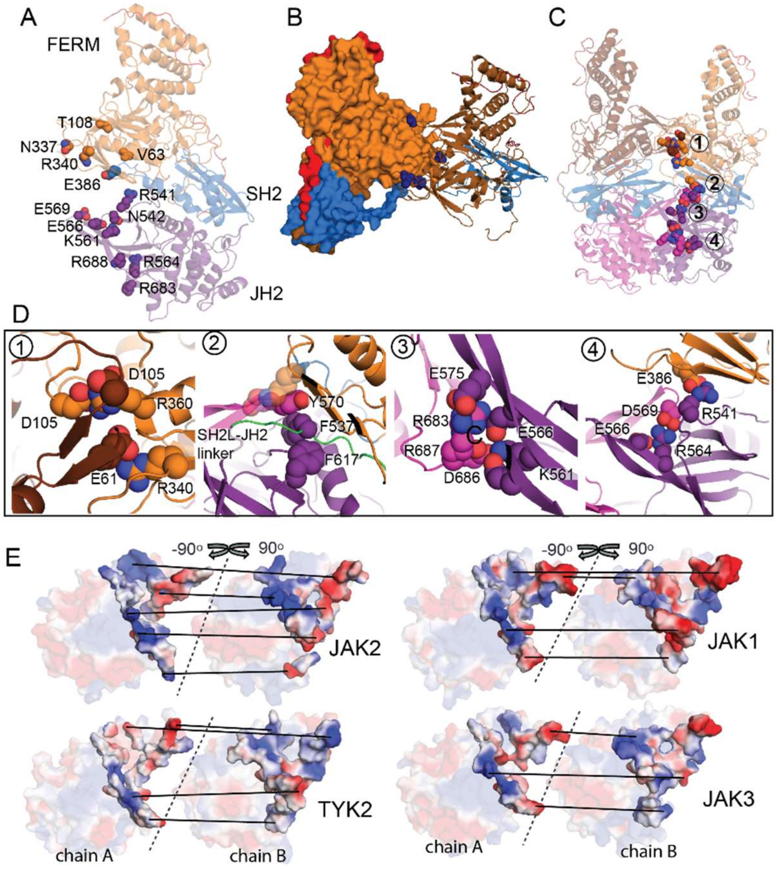


**Figure S3. Related to Figure 3: The active dimer model.** (A) Somatic mutation sites at the dimer interface. (B) Somatic mutation sites residing at the FERM-FERM interface (dark blue spheres). (C) Key residues at the dimer interface. (D) Close-up view of the interface regions numbered in C. (E) Electrostatic features of the JH2-JH2 interfaces in the active homodimer models of JAK2, JAK1, TYK2, and JAK3. Each line connects two matching regions at an interface. Domains are colored as in Figure 1.

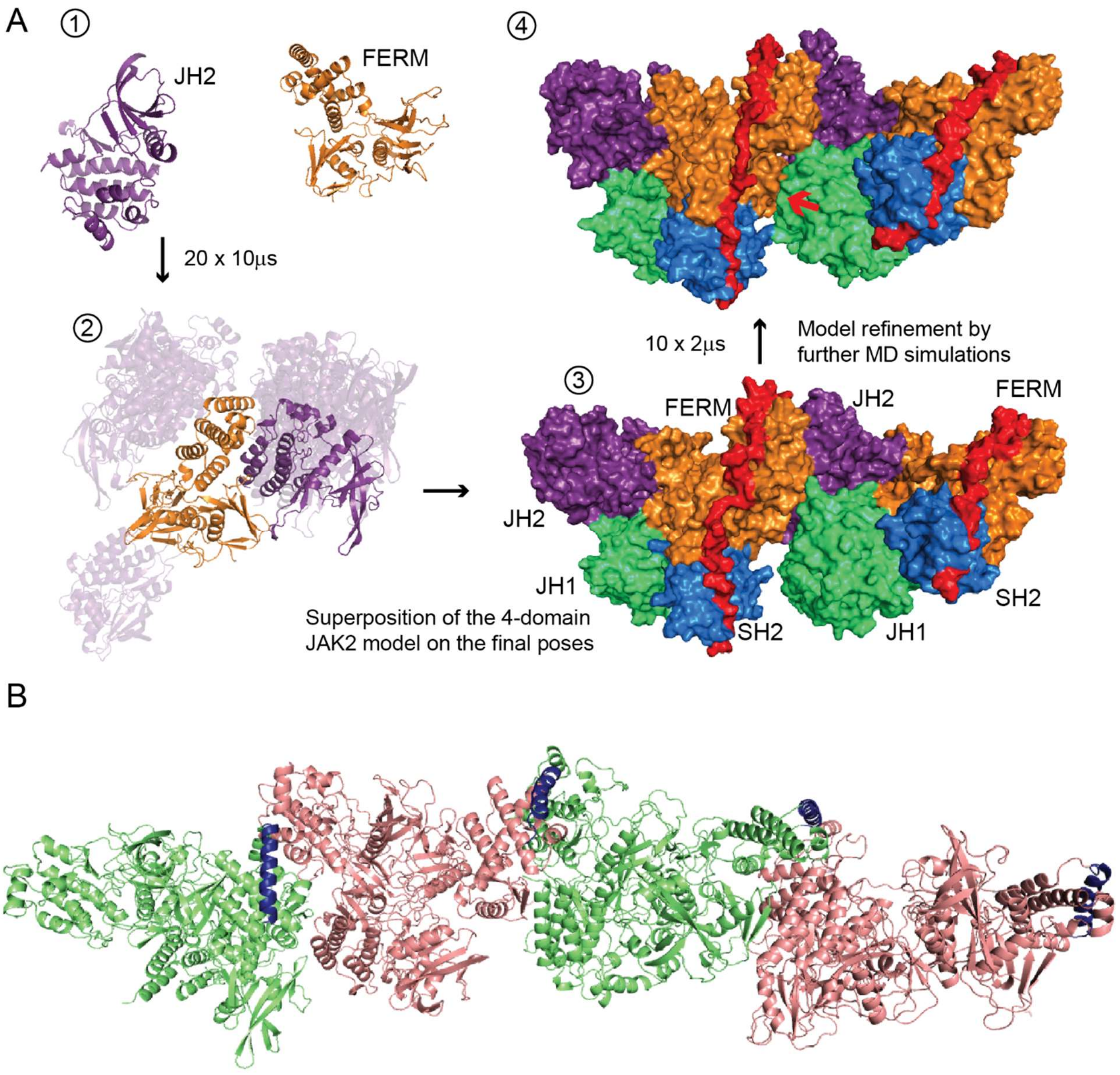


**Figure S4: The speculative inactive dimer model.** (A) Construction of the model: (1) simulations (20 at 10 μs each) of association between FERM and JH2; (2) the 20 FERM-JH2 poses generated by the simulations, with the one we used in subsequent modeling shown in solid colors; (3) two copies of the inactive JAK2 monomer model assembled based on the selected FERM-JH2 pose; (4) the inactive JAK2 dimer model after further simulations. Domains are colored as in Figure 1. (B) The JAK2 oligomer model derived from extending the (asymmetric) inactive dimer model.

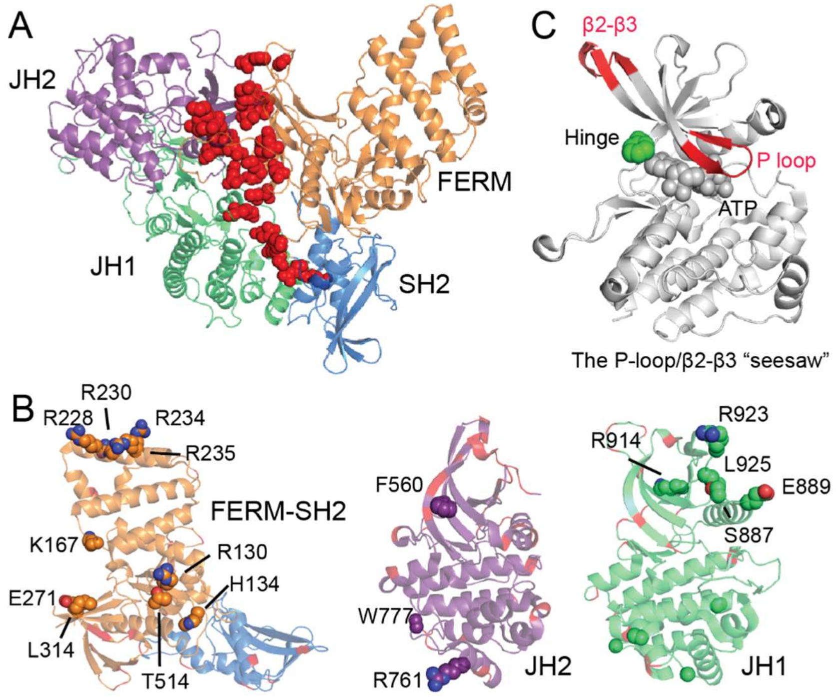


**Figure S5. Related to Figure 4: Somatic cancer mutations.** (A) Somatic mutations at the interface between the FERM-SH2 and JH2-JH1 units (red spheres) in the JAK2 inactive monomer model. (B) Somatic mutations that are not at any interfaces suggested by our models (colored spheres) (see Table S1). Domains in A and B are colored as in Figure 1. (C) The JH2 "seesaw" involving the P-loop and β2-β3 loop.

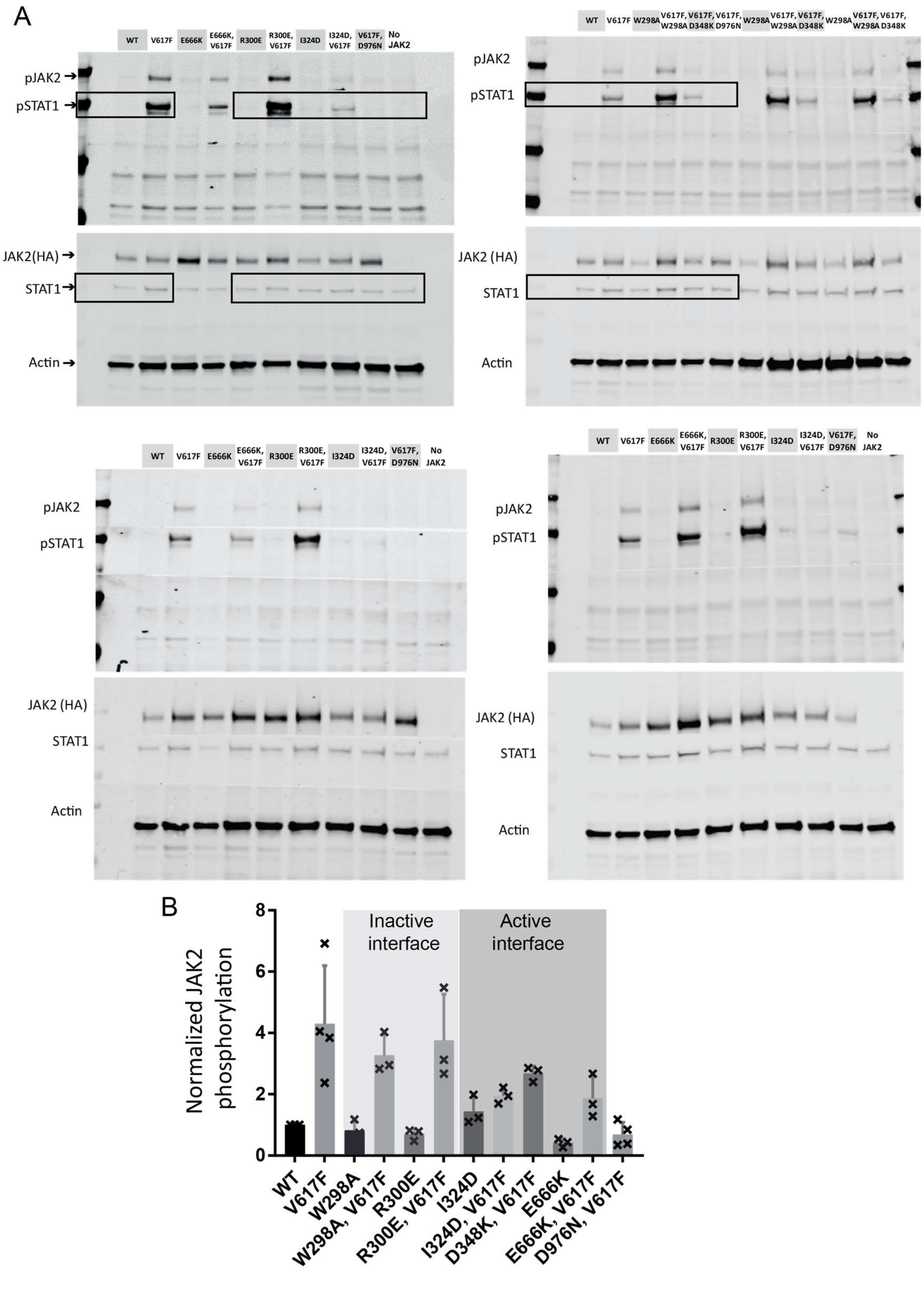


**Figure S6. Related to Figure 5: Western blot data.** (A) Uncropped images of western blots shown in Figure 5B (upper images). For each blot (labeled accordingly), boxes mark the borders of the cropped images shown in Figure 5B. (B) JH1 phosphorylation normalized relative to that of wild-type JAK2. Average values and standard deviations are from at least three independent experiments (n = 3, 4); each x denotes an individual value.

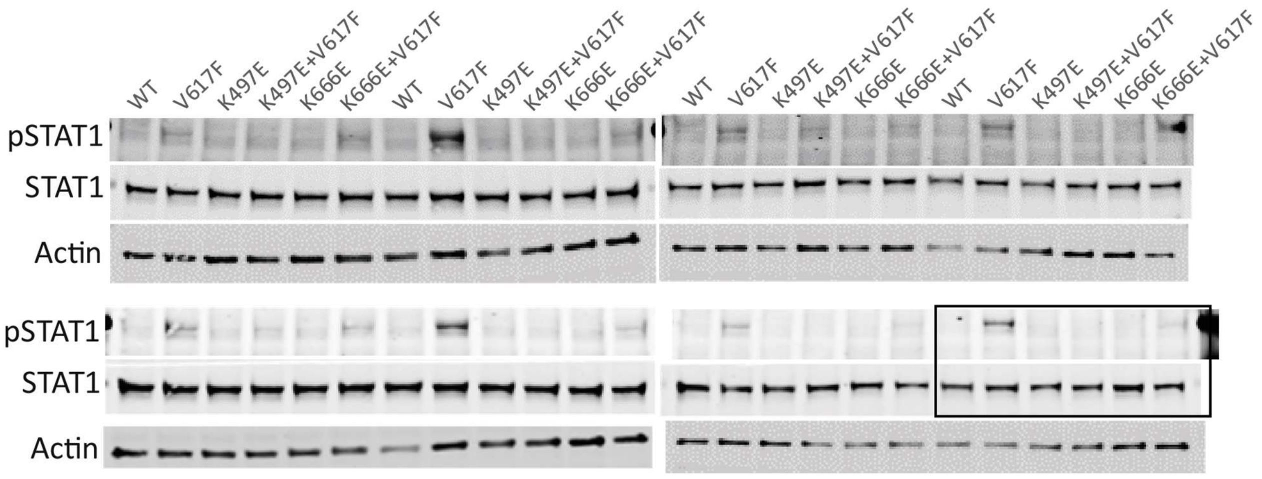


**Figure S7. Related to Figure 5: Additional western blot data.** Uncropped images of western blots shown in Figure 5B (lower image). For each blot (labeled accordingly), boxes mark the borders of the cropped images shown in Figure 5B.
